# A comprehensive update to the *Mycobacterium tuberculosis* H37Rv reference genome

**DOI:** 10.1101/2022.07.15.500236

**Authors:** Poonam Chitale, Alexander D. Lemenze, Emily C. Fogarty, Avi Shah, Courtney Grady, Aubrey R. Odom-Mabey, W. Evan Johnson, Jason H. Yang, A. Murat Eren, Roland Brosch, Pradeep Kumar, David Alland

**Affiliations:** Division of Infectious Disease, Department of Medicine and the Ray V. Lourenco Center for the Study of Emerging and Re-emerging Pathogens, and Rutgers University – New Jersey Medical School, Newark, New Jersey, USA; Public Health Research Institute, Rutgers University – New Jersey Medical School, Newark, New Jersey, USA; Department of Pathology, Immunology and Laboratory Medicine, New Jersey Medical School, Rutgers—The State University of New Jersey, Newark, NJ, USA; Department of Medicine, University of Chicago, Chicago, IL, USA; Committee on Microbiology, University of Chicago, Chicago, IL, USA; Department of Microbiology, Biochemistry and Molecular Genetics, Rutgers University- New Jersey Medical School, Newark, New Jersey, USA; Helmholtz Institute for Functional Marine Biodiversity (HIFMB), Oldenburg, Germany; Division of Computational Biomedicine, Boston University School of Medicine and Bioinformatics Program, Boston University, Boston, MA, USA; Bioinformatics Program, Boston University, Boston, MA, USA; Institut Pasteur, Unit for Integrated Mycobacterial Pathogenomics, CNRS UMR 3525, Paris, 75015, France

## Abstract

H37Rv is the most widely used *M. tuberculosis* strain. Its genome is globally used as the *M. tuberculosis* reference sequence. We developed Bact-Builder, a pipeline that leverages consensus building to generate complete and highly accurate gap-closed bacterial genomes and applied it to three independently sequenced cultures of a parental H37Rv laboratory stock. Two of the 4,417,942 base-pair long H37Rv assemblies were 100% identical, with the third differing by a single nucleotide. Compared to the existing H37Rv reference, the new sequence contained approximately 6.4 kb additional base pairs encoding ten new regions. These regions included insertions in PE/PPE genes and new paralogs of *esxN* and *esxJ*, which were differentially expressed compared to the reference genes. Additional sequencing and assembly with Bact-Builder confirmed that all 10 regions were also present in widely accepted strains of H37Rv: NR123 and TMC102. Bact-builder shows promise as an improved method to perform extremely accurate and reproducible *de novo* assemblies of bacterial genomes. Furthermore, our findings provide important updates to the primary tuberculosis reference genome.

## Introduction

*Mycobacterium tuberculosis* is estimated to infect roughly a quarter of the world’s population and was the second leading infectious disease killer after Sars-CoV2 in 2020 ^1^. The first *M. tuberculosis* genome was sequenced in 1998 by Cole et. al ^2^. They described the complete genome of the *M. tuberculosis* strain H37Rv ^2^, which was first isolated from a patient with pulmonary tuberculosis in 1905^2–4^. Although the original strain has been lost, strains ATCC 27294 (TMC 102) and ATCC 25618 (NR-123) were both isolated from the same patient in separate years and are frequently used as H37Rv in studies^5^. H37Rv remains the most widely used *M. tuberculosis* strain for laboratory experimentation, and the 1998 H37Rv whole genome sequence is accepted as the *M. tuberculosis* reference sequence.

Since then, next-generation sequencing (NGS) tools have been used to generate short reads (75-300 bp) that can be rapidly *de novo* assembled to create draft genomes. However, these assemblies are typically fragmented, harbor mapping artifacts, and contain misannotated gene calls^6^. This problem is largely addressed by using a reference-based assembly approach which maps reads to a known and extensively curated reference genome such as H37Rv. Despite its widespread use, uncertainties remain regarding the completeness of the currently available H37Rv sequence. Repetitive sequences, gene duplications and gene inversions are known confounders of whole genome assembly tools^7^, and *M. tuberculosis* is particularly subject to these limitations given that approximately 10% of the *M. tuberculosis* genome is comprised of highly repetitive genes ^8^. Known differences among different H37Rv isolates further confounds this issue ^5^. These findings suggest that a reanalysis of the *M. tuberculosis* H37Rv reference using more accurate genome assembly tools might reveal important new elements of this genome that could impact all studies that use H37Rv as the genome reference.

The development of third-generation sequencing tools such as single molecule sequencing technology that produces (1 kb - >100 kb) long reads enable successful resolution of repetitive or gapped regions and help generate complete and closed assemblies of microbial genomes ^7,9,10^. These new technologies can substantially improve the accuracy of reference genomes and can enable direct comparisons among different bacterial genomes without first performing a reference-based assembly. Direct genome comparisons can be particularly useful when organisms contain “accessory” sequences that are not present in a standard reference genome. By definition, accessory sequences cannot be mapped to a reference, and this results in their exclusion from any reference-based analysis. Several tools have been designed to *de novo* assemble NGS and third-generation sequencing data ^11–17^; however, the variety of these tools and algorithms lead to inherent differences in their outputs^3,11,13,14^, and there is no standard approach for *de novo* bacterial genome assembly. Existing programs are largely limited by a long- or short-read only approach, and/or the use of a specific single program to assemble the genome ^18–20^. Hybrid assembly tools bridge this gap but typically use short read sequencing to build an initial scaffold followed by long reads on top to create a draft genome ^17,21^. These approaches often ignore the shortcomings of individual assemblers and those that rely on a long read only approach must contend with the large number of SNPs and small indels inherently present in long read sequencing and not caught by older basecalling tools. Furthermore, very few programs or pipelines exist that enable end-to-end generation of highly accurate gap-closed genomes starting with raw sequencing data.

In our efforts to readdress the *M. tuberculosis* H37Rv sequence and support direct comparisons among *de novo* assembled genomes of clinical *M. tuberculosis* strains, we developed a new pipeline that allows us to generate highly accurate complete whole genome sequences using a *de novo* assembly approach. Named Bact-Builder, this tool uses a long-read consensus-based assembly approach, which is then followed by long and short read polishing. Here, we first demonstrate the virtual 100% accuracy and reproducibility of this approach when sequencing three separate H37Rv cultures as well as the ability of the assembly tool to reveal ∼6.4 kb of new sequence that is absent from in the GenBank (NC_000962.3) published reference, including 10 major regions of difference. Together, this work reveals an important update to the H37Rv references sequence and shows the utility of Bact-Builder for assembling highly accurate bacterial genomes, both for use as genomic references or applied to other metagenomic and pangenome studies of bacterial species.

## Results

### Building and Validating Bact-Builder

Bact-Builder (Figure 1A) was designed to generate highly accurate gap-closed *de novo* bacterial genomes. Bact-Builder steps were first designed using *in silico* generated reads created with BadReads^22^ and ART ^23^ which removed common experimental variables that can affect sequencing output and quality (see Supplementary Materials). Further testing was then performed by experimentally sequencing three independent cultures of the same H37Rv stock, extracting genomic DNA, preparing sequencing libraries and sequencing each culture separately. Sequencing was performed on both ONT MinION and Illumina NovaSeq (paired end 2×150) platforms, and the raw sequencing data was used to assemble each of the three replicates (H37Rv.1-3) using Bact-Builder (Figure 1A). We first tested the accuracy and reproducibility of four commonly used long read assembly tools, Canu, Flye, Miniasm and Raven run in triplicate for all samples. We found significant differences between assemblers across all three sequenced H37Rv samples (Figure 1B-D; S3 A-B). Unlike the analysis with our *in silico* data, the analysis of the genomes reconstructed from the three *in vitro* sequenced samples using a pan-genomics approach revealed critical differences between the reporting of individual assemblers both within a sample and across multiple samples (Figure 1E; S3 C-D). These differences included missing core genes and spurious accessory genes compared to the reference genome, suggesting that the existing tools to process long read sequencing data are not suitable to reconstruct microbial genomes when accuracy and reproducibility are critical concerns.

**Figure 1.**
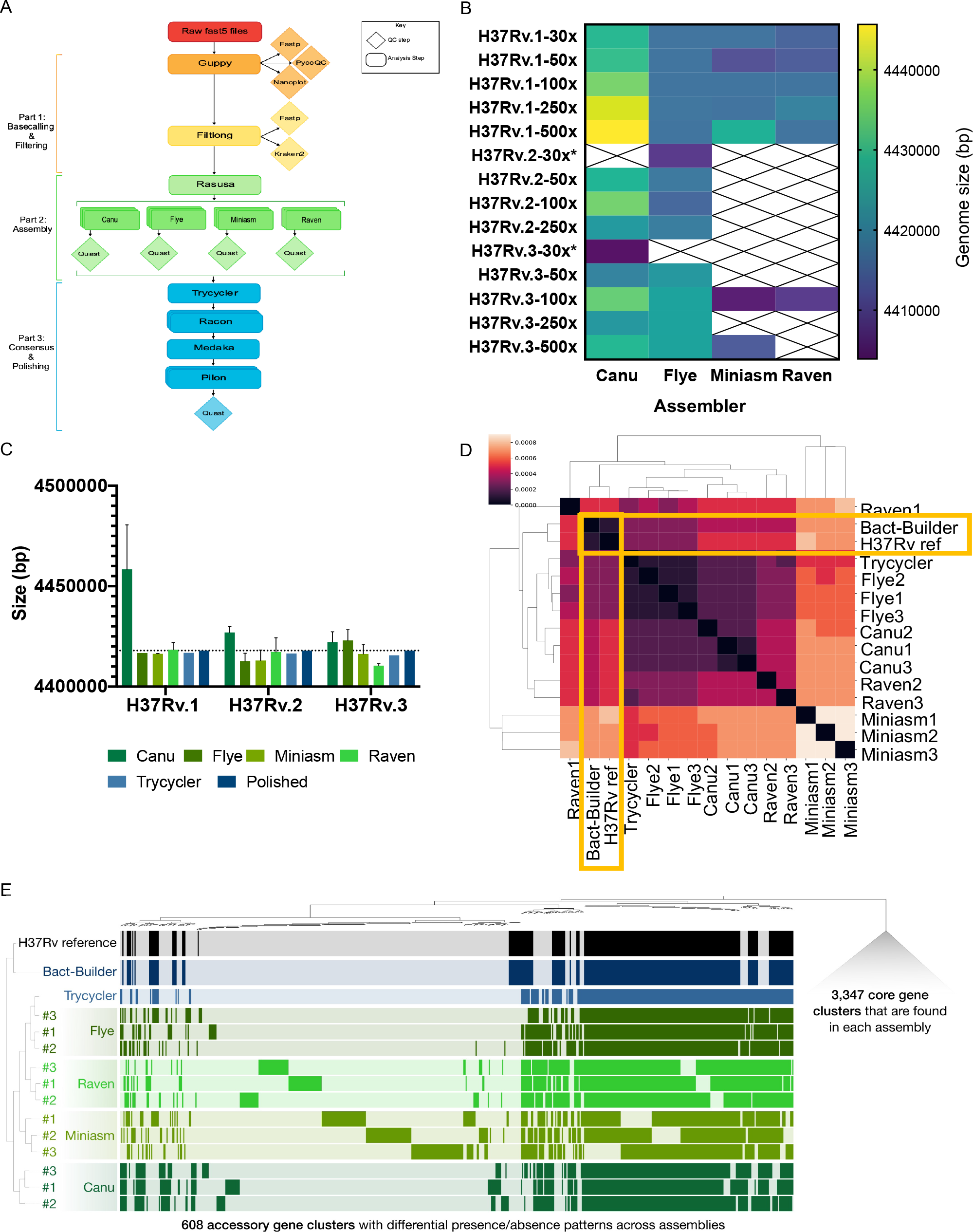
Developing Bact-Builder. **A**. Pipeline overview. Bact-Builder takes raw fast5 sequencing data, filer, assemble, generate a consensus, and polish bacterial genomes. **B**. Heatmap comparison of genome sizes of four *de novo* long read assemblers from laboratory stocks of H37Rv sequenced in triplicate (H37Rv.1-3). The sequence coverage sampled for each analysis is shown in each row on the Y axis. Boxes marked by an X indicate that the assemblies did not pass the trycycler stage because they could not be reconciled with the other assemblies. * Indicates that 3 out of 4 assemblers could not be reconciled, necessitating that trycyler was run with only 1 assembler. **C**. Three replicates of laboratory stocks of H37Rv (H37Rv.1-3), showed variability in size depending on assembler used, and consistent sizes when trycycler was followed by polishing (Bact-Builder output). Dotted line indicates the size of the established H37Rv reference. **D**. Heatmap of hierarchical clustering of the distance using euclidean average linkage clustering of differences between all assemblies for H37Rv.1, the Bact-Builder output and the published reference (H37Rv ref) determined by DNAdiff. **E**. Anvi’o pangenome comparing gene clusters in the reference (H37Rv ref) and H37Rv.1 individual assemblies, trycycler output and the Bact-Builder output.

We worked to reconcile the differences observed across assemblers using Trycycler ^24^ a program that generates a consensus sequence from the outputs of the four individual assemblers. Trycycler produced contigs that were more similar to the reference than any individual assembly and further polishing with both long and short reads generated even more accurate assemblies (Figure 1C), although some remained too fragmented or could not be reconciled and were dropped by the program (Figure 1B). Examining the contribution of sequence coverage depth to the quality of the final Bact-Builder result, we compared Bact-Builder outputs to the H37Rv.1-3 samples using data subsetted to 30x, 50x 100x, 250x, and 500x average read depth. Unlike *in silico* generated reads (Supplementary figure 1B), we found that 30x coverage did not generate a consensus assembly for 2 replicates (H37Rv.2, H37Rv.3) because 3 out of 4 assemblers had could not be reconciled and had to be dropped (Figure 1B). At 50x coverage and above, we did not observe significant variations in genome size (Figure 1B). Using the >50x coverage Dnadiff indicated that Bact-Builder’s assembly of the three independent H37Rv samples produced were identical in size and whole genome sequence (No SNPs were found between H37Rv.1, H37Rv.2 and H37Rv.3) except for one sample (H37Rv.1) that was one nucleotide shorter than the other two (Table 2). The complete genome of H37Rv.1 is reported here as H37Rv (new) and the full genome sequence has been deposited (See supplementary data files; SRA accession: PRJNA836783).

**Table 1.** H37Rv strains used in this study

| **Table 1.** H37Rv strains used in this study |  |  |  |
| --- | --- | --- | --- |
| **Name** | **Strain ID** | **Source** | **Notes** |
| H37Rv.1-3 |  | Laboratory stock,<br>reportedly from the ATCC |  |
| NR-123 | ATCC 25618<br>NR-123<br>(BEI) | AG Karlson, Trudeau<br>Mycobacterial Culture<br>Collection (TMC) | Derived from E.R. Baldwin's human-lung isolate H37<br>by W. Steenken |
| TMC 102 | ATCC 27294 | GP Kubica, Dissociated in<br>1934 | Derived from E.R. Baldwin's human-lung isolate H37<br>by W. Steenken |
| H37Rv1998 |  | Pasteur Institute | H37Rv stock used to generate reference sequence |

**Table 2.** H37Rv.1-3 individual assembly results, and DNAdiff analysis comparing individual assemblies to the H37Rv reference (H37Rv ref).

| Name | # of contigs | Size (bp) | SNP* count (relative to ref) | Indel** count (relative to ref) | Regions of difference (relative to reference) |
| --- | --- | --- | --- | --- | --- |
| H37Rv ref | 1 | 4411532 | - | - | - |
| H37Rv.1 canu1 | 1 | 4446966 | 109 | 2005 | 10 |
| H37Rv.1 canu 2 | 3 | 4483900 | 110 | 1989 | 13 |
| H37Rv.1 canu 3 | 1 | 4444289 | 110 | 2019 | 10 |
| H37Rv.1 flye 1 | 1 | 4416736 | 123 | 1251 | 9 |
| H37Rv.1 flye 2 | 1 | 4416759 | 133 | 1234 | 9 |
| H37Rv.1 flye 3 | 1 | 4416479 | 126 | 1248 | 9 |
| H37Rv.1 miniasm 1 | 1 | 4416466 | 241 | 2961 | 9 |
| H37Rv.1 miniasm 2 | 1 | 4416251 | 240 | 2841 | 9 |
| H37Rv.1 miniasm 3 | 1 | 4416359 | 240 | 2841 | 9 |
| H37Rv.1 raven 1 | 1 | 4414975 | 159 | 1909 | 11 |
| H37Rv.1 raven 2 | 1 | 4422017 | 140 | 1916 | 10 |
| H37Rv.1 raven 3 | 1 | 4418092 | 149 | 1835 | 11 |
| H37Rv.1 Tricycler | 1 | 4416834 | 110 | 1157 | 8 |
| **H37Rv.1 Bact-Builder (polished)** | **1** | **4417941** | **109** | **36** | **10** |
| H37Rv.2 canu1 | 1 | 4429379 | 113 | 2301 | 9 |
| H37Rv.2 canu 2 | 1 | 4427845 | 111 | 2323 | 9 |
| H37Rv.2 canu 3 | 1 | 4423645 | 110 | 2326 | 9 |
| H37Rv.2 flye 1 | 1 | 4408125 | 125 | 1386 | 10 |
| H37Rv.2 flye 2 | 1 | 4414937 | 133 | 1377 | 9 |
| H37Rv.2 flye 3 | 1 | 4414949 | 137 | 1377 | 9 |
| H37Rv.2 miniasm 1 | 5 | 4406947 | 137 | 1896 | 11 |
| H37Rv.2 miniasm 2 | 4 | 4415501 | 130 | 1246 | 12 |
| H37Rv.2 miniasm 3 | 4 | 4416498 | 125 | 1852 | 10 |
| H37Rv.2 raven 1 | 2 | 4424323 | 107 | 1727 | 24 |
| H37Rv.2 raven 2 | 1 | 4410082 | 197 | 1638 | 30 |
| H37Rv.2 raven 3 | 1 | 4410082 | 197 | 1638 | 30 |
| H37Rv.2 Tricycler | 1 | 4416448 | 114 | 1538 | 8 |
| **H37Rv.2 Bact-Builder (polished)** | **1** | **4417942** | **109** | **35** | **10** |
| H37Rv.3 canu 1 | 1 | 4428104 | 123 | 5436 | 10 |
| H37Rv.3 canu 2 | 1 | 4419097 | 121 | 5537 | 10 |
| H37Rv.3 canu 3 | 1 | 4419406 | 131 | 5465 | 9 |
| H37Rv.3 flye 1 | 4 | 4416882 | 201 | 2825 | 11 |
| H37Rv.3 flye 2 | 4 | 4425435 | 190 | 2846 | 15 |
| H37Rv.3 flye 3 | 4 | 4426660 | 196 | 2885 | 16 |
| H37Rv.3 miniasm 1 | 1 | 4421693 | 158 | 3350 | 11 |
| H37Rv.3 miniasm 2 | 1 | 4414787 | 173 | 3619 | 10 |
| H37Rv.3 miniasm 3 | 1 | 4412379 | 190 | 3609 | 12 |
| H37Rv.3 raven 1 | 1 | 4410598 | 164 | 3385 | 25 |
| H37Rv.3 raven 2 | 1 | 4411303 | 141 | 3134 | 21 |
| H37Rv.3 raven 3 | 1 | 4409434 | 188 | 3119 | 21 |
| H37Rv.3 Trycycler | 1 | 4415585 | 114 | 2400 | 8 |
| **H37Rv.3 Bact-Builder (polished)** | **1** | **4417942** | **109** | **37** | **10** |
| *SNP: Single Nucleotide Polymorphism; ** Indels: single base insertions or deletions |  |  |  |  |  |

### Corrections to the H37Rv genome

A comparison between H37Rv(new) and the H37Rv1998 reference strain genome (NC_000962.3) revealed ten regions (R) of difference (R1 – R10) found only in H37Rv(new) (Figure 2). R1 was a 207 bp in-frame insertion in *PE_PGRS27* (*rv1450c*). R2 was a 1356 bp sequence that contained duplications of *rv3475* and *rv3474*, which together make up an IS6110 transposase. There are 7 copies of this region found in the H37Rv1998 and 8 found in H37Rv.1-3. In R3 a single complete copy of *PPE38* was replaced by an 845 bp sequence that contained an upstream copy of PPE38 and a downstream second truncated copy of *PPE38* (which we have named *PPE38a* or *rv2352c*.2). Between these two PPE38 genes we found a new paralog of *esxN* (*rv1793*) which we have named *esxN*.*2* (*rv1793*.*2*), and a new paralog of the *esxJ* (*rv1038c*) *and esxM* genes (*rv1792*) *(esxM is also called esxJ in M. tuberculosis CDC1551*) ^8,25^, which we have named *esxJ*.3 (*rv1038c*.*2*). The R4, R5 and R6 sequences each contained increased numbers of tandem duplications in intergenic regions (Table 4). R7 was a 1728 bp in-frame insertion at the 3’ end of *PPE54* (*rv3343c*), substantially increasing its size. R8 was a 9 bp insertion of a tandem duplication in *PE_PGRS51* (*rv3367*). R9 was a 579 bp in-frame insertion in the middle of *PE_PGRS54* (*rv3508*), which also significantly increasing the size of the gene. R10 was a 111 bp in-frame insertion in the middle of *PE_PGRS57* (*rv3514*). In addition to the 10 regions, 109 SNPs and 37 indels were found between H37Rv(new) and the H37Rv reference.

**Figure 2.**
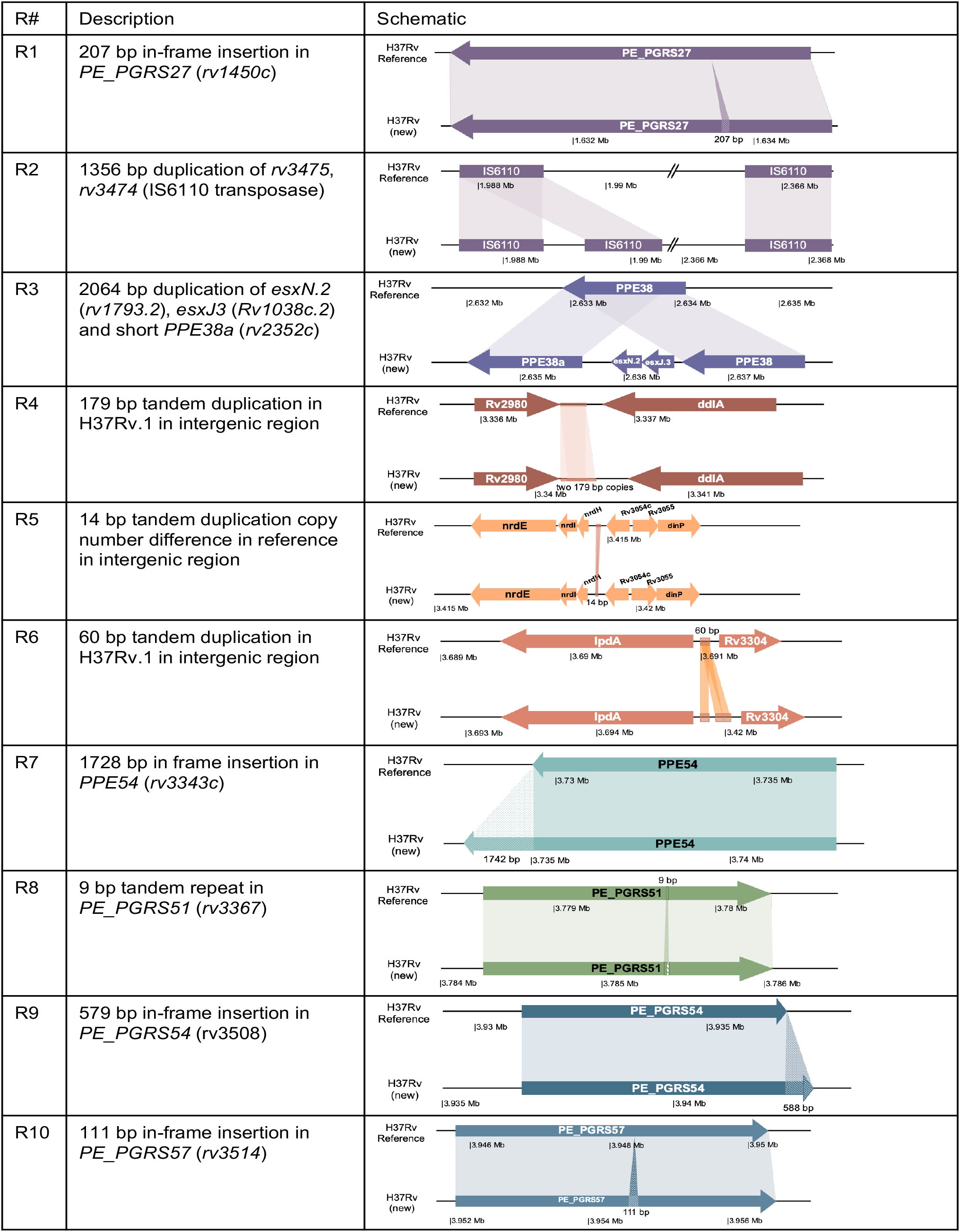
Identifying and validating Regions of difference. Description of the 10 regions of difference identified by DNAdiff between the H37Rv reference and H37Rv (new).

We evaluated the potential functional impact of amino acid differences between *esxN* and *esxN*.*2* or between *esxJ*, (*CDC551-esxJ/esxM*) and *esxJ*.*3* using PROVEAN^26^, which suggested that the paralogs maintained an overall conservation of protein structure (Figure 4A). PCR and subsequent Sanger sequencing of R3 revealed that the observed differences were real and were likely missed using the original shot-gun sequencing approach (Figure 4B). We also PCR amplified H37Rv strains NR123, TMC102, TMC301 and H37Rv1998 and detected R3 in all three strains (Figure 4B). These results are also supported by previous studies which identified the genes found in our H37Rv R3 region in another strain of *M. tuberculosis and Mycobacterium marinum* and *Mycobacterium microti*, although using a different nomenclature^25,27–29^.

Finally, we completely sequenced and used Bact-Builder to assemble NR123, TMC102, and 1998H37Rv. We again detected all 10 regions of difference confirming that R1–10 are true regions shared by all publicly available H37Rv isolates. A phylogenetic analysis of these strains demonstrated that H37Rv.1–3 were most closely related to TMC102, differing by only 4 SNPs (Figure 3). We found that H37Rv1998 was most closely related to NR123, but NR123 still differed from H37Rv1998 by in-frame insertions in *rv2680* (conserved hypothetical protein), *rv3367* (*PE_PGRS51*) and intergenic regions, and by in-frame deletions in *rv0297* (*PE_PGRS5*), *rv3514* (*PE_PGRS57*) and in intergenic regions (Table 3, Figure 3). Compared to NR123, TMC102 and H37Rv.1-.3 had two additional IS6110 transposon elements, and in-frame insertions in *rv0297* (*PE_PGRS5*), *rv209*0 (Probable 5’-3’ exonuclease), *rv3367* (*PE_PGRS51*) and in intergenic regions (Table 3).

**Figure 3.**
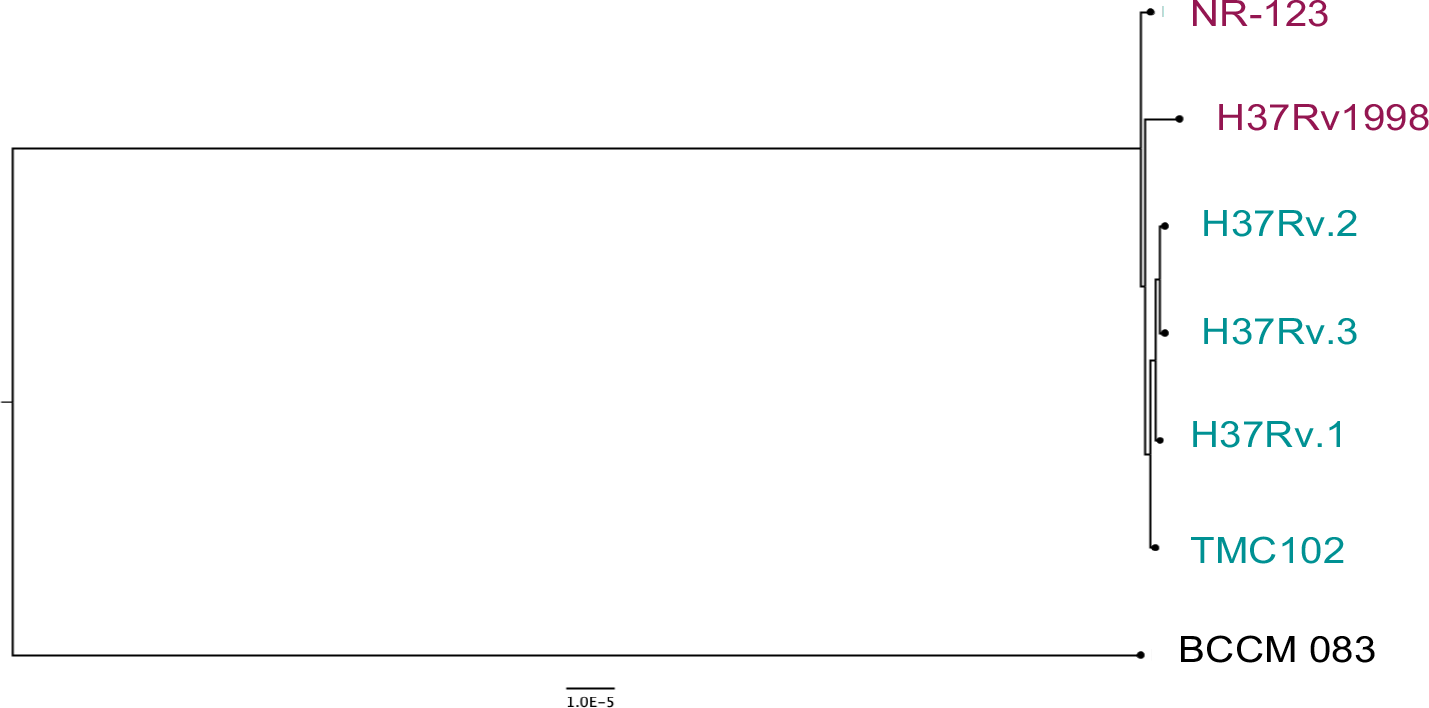
Phylogenetic analysis of H37Rv strains used in the study. H37Rv strains were assembled with Bact-Builder, sequence alignment was completed using Bowtie2 via REALPHY ^67^ (https://realphy.unibas.ch/fcgi/realphy), and a maximum likelihood tree was constructed with RaxML^68^. The tree was rooted with BCCM 083, a M. tuberculosis Lineage 1 strain ^69^.

**Table 3.**
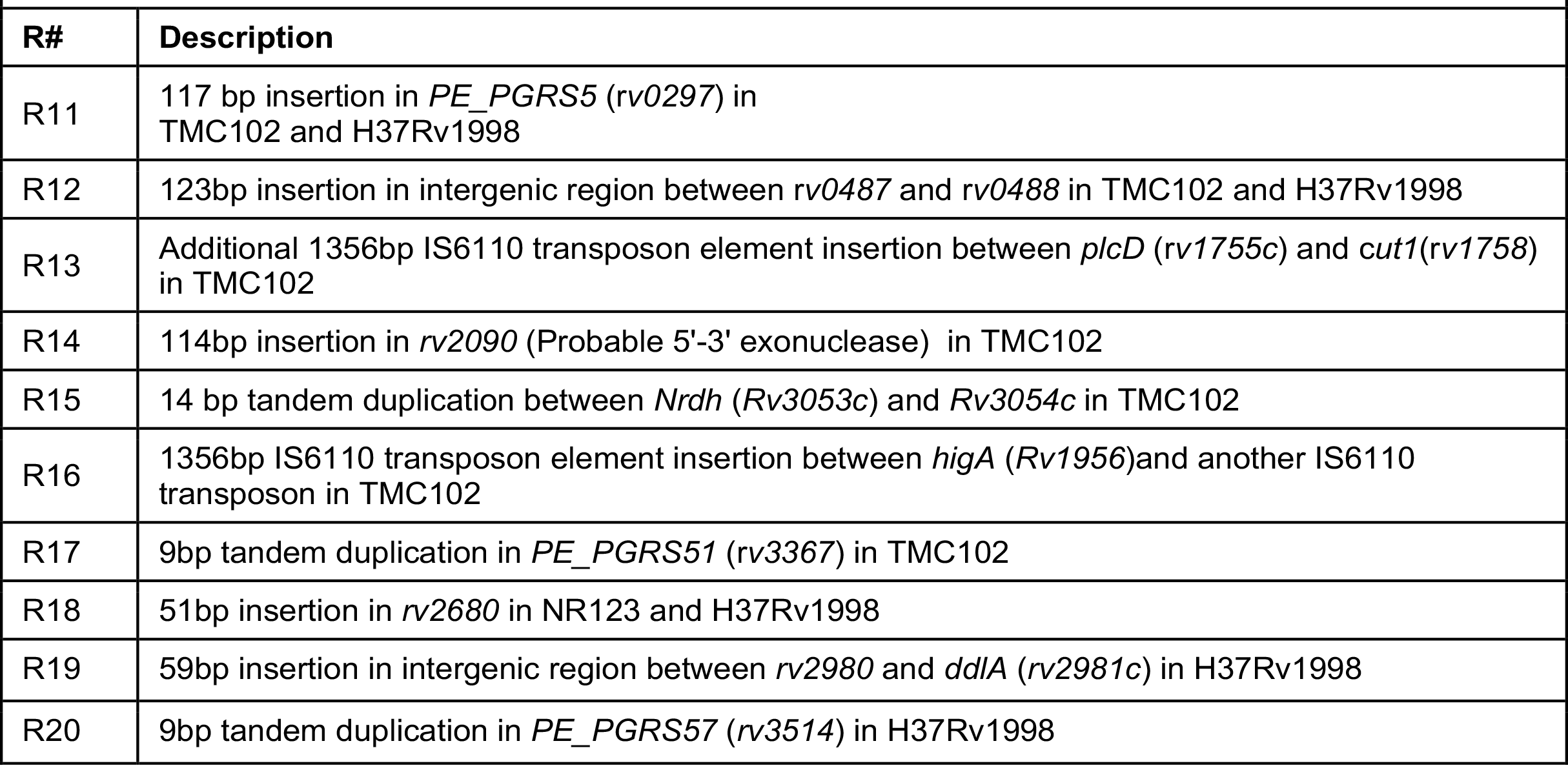
Description of differences between the newly sequenced and *de novo* assembled H37Rv strains, TMC102, NR123 and H37Rv1988 identified by DNAdiff

### All 10 H37Rv Regions are expressed

As further confirmation that none of the regions found in H37Rv(new) were caused by sequencing or assembly artifacts, we tested whether published H37Rv RNA sequencing studies contained reads that mapped to each region. This investigation also allowed us to determine whether the newly discovered *esxN*.*2* and *esxJ*.*3* genes were expressed differently comparted to their *esxN* and *esxJ* paralogs as one possible indication of significant functional differences between these genes. We downloaded all public *M. tuberculosis* H37Rv RNA sequencing datasets on the NCBI SRA database. Of the 908 datasets available at the time of our analysis, 905 datasets used Illumina sequencing chemistry, passed our quality control metrics, and were therefore included in these analyses. We aligned these raw reads to both the existing H37Rv1998 reference genome and to H37Rv(new) and computed the feature counts for each genomic region of difference following batch correction. We discovered that raw RNA reads from public data sets did indeed map onto the newly discovered Regions in H37Rv(new) (Figure S4). However, both *esxN*.2 and *esxJ*.3 showed significantly different levels of expression across the 905 datasets compared to their *esxN* and *esxJ* paralogs (Figure 4C-D). Expression levels between paralogs of *esxN* and *esxJ* respectively were moderately correlated, possibly indicating that the paralogs were under different regulatory control (Figure 4 E, F). Expression between *esxN*.*2* and *esxJ*.*3* however were strongly correlated, likely because the genes are adjacent to each other (Figure 4G).

**Figure 4.**
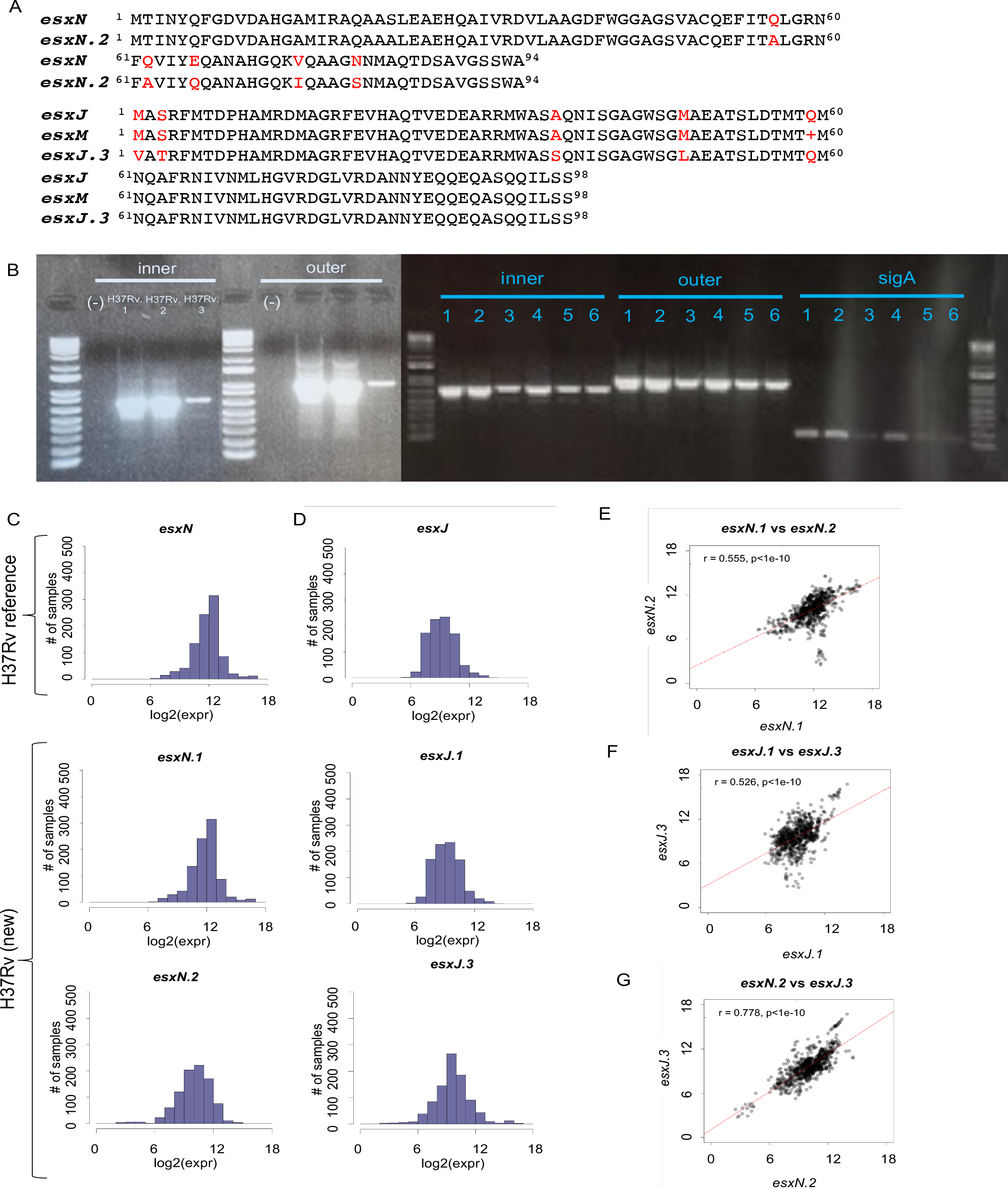
Identifying R3 in all H37Rv strains. **A**. Clustal Omega comparison of amino acid sequence for *esxN* and *esxN*.*2* and between *esxJ, esxM* and *esxJ*.*3*. **B**. PCR of R3 in all 3 laboratory replicates of H37Rv (H37Rv.1-3), H37Rv1998 (1-3) and 3 commercially available strains of H37Rv from ATCC (4:NR-123, 5: TMC102, 6: TMC301). Inner primers targeted inside the region (738 bp) and outer primers targeted flanking regions (1007 bp). *SigA* primers were used as a housekeeping control. **C-D**. Histogram of *EsxN* (**C**) and *esxJ* (**D**) expression in public H37Rv datasets, demonstrating that newly identified *esxN*.*2* and *esxJ*.*3* are expressed in H37Rv and exhibit differential gene expression compared to their respective paralogs. **E-G**. Scatterplots showing correlation of expression in H37Rv (new) of *esxN*.*1* and *esxN*.*2* (**E**); *esxJ*.*1* and *esxJ*.*3* (**F**); and *esxN*.*2* and *esxJ*.*3* (**G**).

## Discussion

The development of new NGS tools and pipelines to generate accurate complete genomes is vital to our understanding of drug resistance evolution, pathogenesis, and virulence of clinically relevant pathogens such as *M. tuberculosis*^44^. The first complete whole genome sequence of the H37Rv strain generated using shot gun sequencing was published in 1998 ^2^ and ushered in a revolution in *M. tuberculosis* research. The published sequence led to the discovery of novel protein families responsible for fatty acid and polyketide biosynthesis, drug efflux pumps, PE/PPE proteins, and transposon elements ^2,5,30^. Minor divergences among subcultures of these strains leading to differences in SNPs, indels and transposable elements have also been described ^5^. However, our current study indicates that there are substantial regions of difference between the Genbank H37Rv reference sequence and what is likely to be the true complete genome sequence of H37Rv isolates including NR-123, TMC 102 and a culture of the original 1998 isolate (Table 3). H37Rv is the workhorse of *M. tuberculosis* research and its Genbank sequence is widely used for aligning and analyzing DNA, RNA, and Tn-seq studies. Thus, the importance of establishing a fully accurate reference genome for this strain cannot be overstated. Our *de novo* assemblies led to the discovery of new *esxN* and *esxJ* paralogs in H37Rv.

These genes are members of the conserved ESX-5 locus. The *esxN* (*rv1793*) gene is a member of the *M. tuberculosis* 9.9 subfamily and is a paralog of *esxA* (ESAT-6). The *esxJ* (*rv1038c*) gene is a member of the QILSS subfamily and is a paralog of *esxB* (CFP-10) ^31^. Canonical *esxA* and *esxB* proteins interact to form a heterodimer that is secreted from *M. tuberculosis*. These proteins are known to be frequently recognized T-cell antigens ^32^. Several *esx* gene pairs are contained within the conserved ESX-1 – ESX-5 loci which together encompass a type VII secretion system (T7SS). ESX-5 has been linked to PE and PPE protein secretion during macrophage infection ^33,34^. Loss of ESX-1 has been linked to attenuation in *M. bovis* BCG ^35^ and there is evidence for attenuation in ESX-5 mutants in *M. tuberculosis* as well ^36^. Not only are the new ESX-5 genes which we identified actively transcribed, but they are also differentially expressed compared to the known ESX-5 genes. This strongly suggests that *esxN2* and *esxJ3* may have some new functionalities compared to their previously paralogs, including a potential a role in *M. tuberculosis* pathogenesis. Regions with similar genomic organization to R3 were previously observed in *M. marinum* ^27^, *M. microti* ^29^ and *M. tuberculosis* strain CDC1551^27^. However, our findings suggest that the genes previously annotated as *esxX, esxY* and *PPE71* should instead be named *esxN*.*2, esxJ*.*3* and *PPE38* based on sequence homology and PGAP annotation of our fully closed genome. The *esxJ* gene in the H37Rv reference sequence and our newly annotated *esxJ*.*3* also share partial sequence homology to *rv1792* which is annotated as *esxM* in the reference genome due to the occurrence of a stop codon in the middle of the gene, substantially shortening the overall protein size. Additionally, *esxM* was correctly annotated as *esxJ* in the *M. tuberculosis* strain CDC1551 annotation despite presence of the stop codon. We have therefore named the new paralog discovered in the H37Rv as *esxJ*.*3* to avoid confusion, even though there is no esxJ.2 in the H37Rv reference.

The *M. tuberculosis* PE/PPE genes are a group of approximately 170 genes that are named for the presence of N-terminal ProGlu and ProProGlu motifs. These genes account for roughly 10% of the *M. tuberculosis* genome ^2,37,38^. PE/PPE genes represent a diverse set of genes that have been implicated in *M. tuberculosis* infection, host immune modulation, nutrient transport ^28,37,38^. We discovered substantial differences in five known PE/PPE genes including the identification of a new PPE gene (PPE38a). The functional implications of our findings are unknown but further investigation will be made possible and feasible with the availability of the newly complete and correct sequences

As prevalence of whole genome sequencing (WGS) for studying large communities of related bacteria increases, accurate reference genomes and methods for *de novo* whole genome assembly will be needed to avoid sequencing artifacts. Our work demonstrates the importance of using a *de novo*, consensus assembly approach for generating high quality genomes. These complete and highly accurate assemblies can enhance our understanding of genomic variation and its role in microbial pathogenesis and evolution. In the case of *M. tuberculosis*, our approach provides several improved and complete reference genomes for the second largest cause of death from an infectious disease after COVID-19. The updated genomes of TMC102, NR123 and H37Rv1998 are quite similar as they all contain R1–R10. However, we did note small differences between each isolate and have deposited updated sequences for each (SRA accession#: PRJNA836783). TMC102 appears to be used more commonly used by the global scientific community than NR123, being cited >1000 times versus 246 ^39^ respectively at the time of this writing, suggesting that TMC102 should perhaps become the single accepted laboratory strain. Regardless of which H37Rv strain is used in any given experiment, our work suggests that the use of primary ATCC stocks and frequent WGS followed by *de novo* assembly should be performed to ensure that the genome of the H37Rv being used is sufficiently conserved to allow comparisons between research groups. Indeed, our approach can be applied to any individual bacterial genome to provide a complete, closed genome and is an invaluable tool towards development of comprehensive and accurate genomes. Ultimately this will enable the assembly of a new global set of *M. tuberculosis* reference genomes that together describe a true *M. tuberculosis* pangenome and will fuel studies that lead to a better understanding of tuberculosis pathogenesis, strain variation, and ultimately new ways to eliminate this age-old disease.

## Methods

### Bacterial strains

Laboratory stocks of *M. tuberculosis* H37Rv (H37Rv.1-3) were used for all real-strain experiments unless otherwise noted. The H37Rv strain (called H37Rv1998 in this work) was the source of the BAC library^40^ used to generate the original H37Rv genome sequence described in 1998^2^ was obtained from Institute Pasteur, France. Two additional H37Rv strain variants NR-123 (ATCC 25618), and TMC 102 (ATCC 27294), were obtained from BEI resources^41^ and American type culture collection (ATCC)^39^ respectively (Table1).

### Culture and DNA extraction

*M. tuberculosis* strain H37Rv (Table 1) was cultured in Middlebrook 7H9 (BD Difco, Franklin Lakes, USA), 10% oleic acid-albumin-dextrose catalase (OADC) (BD Difco, Frankin Lakes, USA), 0.05% (vol/vol) Tween 80 (Sigma-Aldrich, St. Louis, USA) and 0.2% (vol/vol) glycerol (Sigma-Aldrich, St. Louis, USA). Late exponential phase cultures were plated on 7H11 (Sigma-Aldrich, St. Louis, USA) plates + 0.5% (vol/vol) glycerol and extracted using a slightly modified Cetyltrimethylammonium bromide (CTAB) method ^42^. Briefly, plated strains were grown until confluent (∼3 weeks). Roughly 1/3 of the plate (2-3 loopfuls) were added to 400 µl 1x Tris-Ethylenediaminetetraacetic acid (EDTA) pH 7.5 (Thermo Scientific, Waltham, USA) and heated at 80 ºC for 30 min. Lysozyme (Sigma-Aldrich, St. Louis, USA) was added to the tube (final concentration 20 mg/ml), followed by incubation at 37ºC overnight. Seventy µl of 10% sodium dodecyl sulfate (SDS) (w/v) (Promega, Madison, USA) and proteinase K (final concentration 0.2 mg/ml; 10 mg/ml) (Invitrogen, Waltham, USA) was added and the sample was incubated at 50 ºC for 20 min. A mixture of *N*-acetyl-*N,N,N*-trimethyl ammonium bromide (CTAB; final concentration, 40 mM) and NaCl (final concentration, 0.1 M) was added, followed immediately by the addition of NaCl alone (final concentration, 0.6 M) and the sample was incubated at 50 ºC for 10 min, then 700 µl of chloroform-isoamyl alcohol (24:1) (Sigma-Aldrich, St. Louis, USA) was added. The sample was then pulse vortexed and then centrifuged for 6 min at 16903 xg at room temperature. The upper aqueous phase was transferred to a new tube taking care to avoid removing any part of the interface, then 5 µl of RNAse A (10 mg/ml; Qiagen, Hilden, Germany) was added and the sample was incubated for 30 min at 37 °C. Seven hundred µl of chloroform-isoamyl alcohol (24:1) was then added to the sample for a final extraction. The sample was pulse vortexed and centrifuged for 6 min at 16903 xg at room temperature. The genomic DNA in the resulting aqueous phase was isolated by 1x volume cold isopropanol precipitation. Spooled DNA was removed and washed with 70% cold ethanol. The sample was then spun at 16903 xg for 2 min at 4 °C, the ethanol was removed, and the sample was air dried. The final sample was resuspended in 50 µl nuclease free water.

### Genomic DNA (gDNA) sample quality check

Once extracted, H37Rv gDNA concentration and quality was evaluated using a Nanodrop One™ (ThermoFisher, Waltham, USA). Fragment size and DNA integrity number (DIN score) was evaluated using Genomic DNA Screen Tape on a 4200 TapeStation system (Agilent, Santa Clara, USA). H37Rv gDNA extracts that met the following cutoffs were used for sequencing: A260/280 ratio: between 1.7-2.0; A260/230 ratio: between 1.5-2.5; DIN score ≥ 7; and DNA length > 20 kb. Purified DNA was stored at 4 °C prior to preparation of sequencing libraries.

### Whole genome sequencing (WGS)

*Nanopore Sequencing*. Three independent replicates of H37Rv laboratory stock (H37Rv.1-3) gDNA were processed with a 1D sequencing kit (SQK-LSK109) (Oxford Nanopore Technologies (ONT, Oxford, United Kingdom) along with a native barcoding kit (EXP-NBD103 and EXP-NBD112) according to the native barcoding gDNA protocol (Supplementary table 1). The gDNA was not sheared but used directly for DNA end repair and ligation. Both ligation steps in the protocol were extended from 10 min to 30 min. To ensure enough library was available for each run, the pooled adapter ligation step was performed in duplicate, and the final libraries were pooled before sequencing. The final library was sequenced using a FLO-MIN106 flow cell on a MinION instrument.

### Illumina PCR-free sequencing

WGS Illumina reads were used to polish consensus sequences for all samples included in this study. The H37Rv samples were library prepped using an Illumina DNA PCR-free library prep kit (Illumina, San Diego, USA) (Supplementary table 1). Sequencing libraries were quality checked using the Qubit™ ssDNA kit (Thermo Fisher, Waltham, USA) and equal volumes were pooled for sequencing. The libraries were then sequenced in a paired-end 150 bp configuration on an Illumina NovaSeq 6000 platform.

### Artificial *in silico* read generation

Bact-Builder development and testing made use of artificial reads generated for *M. tuberculosis* H37Rv, the most commonly used laboratory adapted *M. tuberculosis* reference strain. A published H37Rv fasta obtained from NCBI (NC_000962.3) was run through BadRead (v0.1.5; –quantity 100x --length 1000, 1000) (https://github.com/rrwick/Badread) in order to generate simulated ONT reads ^22^. *In silico* paired-end Illumina reads were generated with ART ^23^ using the same published H37Rv fasta and the default ART parameters to generate Illumina 150 bp paired-end reads with a fold coverage of 30x.

### Assembly Pipeline

A full reproducible pipeline encompassing all these steps herein was built within Nextflow ^43^ and can be found at github.com/alemenze/bact-builder. Parameters and software versions listed below are defaults within the automated pipeline. An overview of the entire pipeline can be found in Figure 1A.

### Compute resources

The pipeline was run within the local Amarel HPC environment but has been designed to be optimized to custom HPC or cloud-based resources. The local environment consisted of base nodes with 2x Intel Xeon Gold 6230R (Cascade Lake) Processors (35.75 MB cache, 2.10 GHz): 2933 MHz DDR4 memory, 26-core processors (52 cores/node), 12×16 GB DIMMS (192 GB/node) per node, and GPU nodes included graphics cards with the NVIDIA Pascal architecture.

### Basecalling and demultiplexing

Raw nanopore sequencing reads were base-called and demultiplexed with the latest version of the Guppy basecaller available at the point of sequencing (Guppy v.4.2.2). Sequencing quality and output was evaluated using Nanoplot (v1.33.0) (https://github.com/wdecoster/NanoPlot) ^44^ and pycoQC (v2.5.0.23) (https://github.com/tleonardi/pycoQC)^45^.

### Quality control and filtering

Basecalled and demultiplexed files were trimmed using Filtlong (v0.2.0; --min_length 1000 --min_mean_q 70) (https://github.com/rrwick/Filtlong). Sample contamination was evaluated via taxonomic classification using Kraken2 (v2.1.1) (https://github.com/DerrickWood/kraken2) ^46,47^. Illumina data was trimmed using trim-galore (v0.6.6) (https://github.com/FelixKrueger/TrimGalore)

### Random sub-setting

Nanopore reads were randomly sub-selected to even read depths and variant samples using Rasusa (v0.3) (https://github.com/mbhall88/rasusa).

### Assembly

Nanopore reads for each sample were *de novo* assembled using the following four assemblies in triplicate: Canu (v1.5; genomeSize= 5m)^11^, Flye (v2.8.1-b1676; genome-size 5m --plasmids) ^12^, Miniasm (v0.3-r179) ^48^, and Raven (v1.3.0) ^13^. Gap-closed sequences were evaluated in Bandage (v0.8.1) (https://github.com/rrwick/Bandage) ^49^.

### Consensus assembly

Trycycler (v0.3.0) (https://github.com/rrwick/Trycycler) was used to generate a consensus long-read assembly using the twelve individual assemblies. The cluster step was run with the following modifications: trycycler cluster --min_contig_depth 0.5. Within Bact-Builder, cluster001 is used by default for downstream processes. Trycycler can be run manually for bacteria with extra-chromosomal plasmids or if assemblers need to be removed. Trycycler reconcile was run with the following modifications: trycycler reconcile --cluster_dir trycycler/cluster_001 --max_length_diff 1.3 --max_add_seq 10000 --min_identity 95 --max_indel_size 1000. All other steps were run with default parameters. Following the reconcile step, contigs that could not be reconciled or had a high degree of dissimilarity from the majority of assemblies were removed and the reconcile step was run again. As long as ≥ 6 assemblies passed the reconcile step, trycycler was run to completion and the consensus was used for downstream analysis.

### Polishing

Following trycycler, the consensus assemblies were further polished using both long and short read polishers. Racon (v1.4.20; -m 8 -x -6 -g -8 -w 500) (https://github.com/isovic/racon) was run three times on each assembly using ONT reads. Following racon, medaka (v1.0.1; -m r941_min_high_g360) (https://github.com/nanoporetech/medaka) was used for long read polishing. Finally, Pilon (v1.24; --fix all --changes) (https://github.com/broadinstitute/pilon) was run three times on the medaka consensus using Illumina reads ^50^.

### Evaluating Outputs

Final assemblies were assessed for SNPs, indels and regions of difference using DNAdiff (v1.3) from the MUMmer program (https://github.com/mummer4/mummer) ^51,52^ and quast (v5.0.2). Complete, polished assemblies were also compared to each using a pangenomics approach implemented in anvi’o ^53^ (https://anvio.org) to investigate large structural changes and differences in the accessory gene pool as a function of the assembly and curation steps. Annotation of the consensus genome sequence was performed using the NCBI’s prokaryotic genome annotation pipeline ^54^

### PCR and Sanger sequencing

Following DNAdiff analysis, identified regions were PCR validated by designing PCR primers both within the identified region and flanking regions. PCR primers were designed using Primer Quest (IDT, Coralville, USA). GoTaq Green Master Mix (Promega) was used to run PCR for 30 cycles using manufacturers suggested parameters ^55^. PCR products were run on a 1% Agarose gel containing .08% ethidium bromide and visualized using the ChemiDoc imaging system (BioRad, Hercules, USA). PCR amplicons were sanger sequenced (Psomagen, Rockville, USA) and aligned to the reference to validate the region.

### Expression analysis

Accession numbers for all *M*.*tuberculosis* H37Rv RNA-sequencing experiments publicly available on the NCBI Sequence Read Archive (SRA) as of October 1, 2021 were downloaded using the NCBI’s esearch and efetch utilities, similar to Yoo et al ^56^. Sequencing runs were filtered to include only those performed with Illumina short read sequencing, and runs were processed as described in Ma et al 2021^57^. Sequences were downloaded using their SRA accession numbers using fasterq-dump ^58^. Raw sequence quality was assessed using fastqc ^59^ and adapter trimming was performed using bbduk ^60^. Trimming and sequence removal based on quality was performed using trimmomatic ^61^. Sequences were aligned to either the existing H37Rv reference (NC_000962.3) or the consensus reference reported here by trycycler using Bowtie2 ^62^. Read counts were compiled using featureCounts ^63^. Quality data, adapter and quality trimming statistics, and alignment and counts metrics were compiled and assessed using multiqc ^64^. Of 908 RNA-sequencing runs available from the strain H37Rv, 905 used Illumina sequencing chemistry and passed quality control metrics and were included in the expression analyses reported here. Batch correction was completed by grouping samples by study, identified by the BioSample ID corresponding to a given run’s SRA accession, and performing quantile normalization on raw counts of reads using qsmooth ^65^. Visualization of data was performed in R ^66^.

## Data availability

Data Collection: Whole Genome Sequencing data for the strains used in this study is available from the NCBI Sequence Read Archive via BioProject accession number PRJNA475130. Updated annotation files have been deposited in GenBank, with primary accession code: CP097325 (GenBank upload pending).

## Code availability

Bact-Builder is available at: https://github.com/alemenze/bact-builder**Error! Hyperlink reference not valid**.. Scripts for the RNASeq analysis can be found at: https://github.com/as2654/rna-seq-tb0/tree/main.

## ACKNOWLEDGEMENTS

Research reported in this publication was supported by the National Institute of Allergy and Infectious Diseases of the National Institutes of Health under award numbers U19 AI 111276 and U19 AI 162598. The content is solely the responsibility of the authors and does not necessarily represent the official views of the National Institutes of Health. The authors acknowledge the Genomics Center at Rutgers New Jersey Medical School (https://research.njms.rutgers.edu/genomics/) and the Office of Advanced Research Computing (OARC) at Rutgers, The State University of New Jersey (http://oarc.rutgers.edu) for providing access to the Amarel cluster and associated research computing resources that have contributed to the results reported here.

## Author information

### Contributions

PC, PK and DA designed the study; PC and ADL developed the bioinformatics tools; PC and CAG library prepped and sequenced the samples; PC, ADL, EF and AS performed data analysis; PC, ECF, AME, and AOM contributed to data visualization. PC and DA wrote the manuscript; ADL, ECF, AS, AOM, WEJ, JHY, AME, RB and PK edited the manuscript

