## Supplementary materials and figures for "A comprehensive update to the *Mycobacterium tuberculosis* H37Rv reference genome"

**Supplementary Information**

**Poonam Chitale**, Alexander D. Lemenze, Emily C. Fogarty, Avi Shah, Courtney Grady, Aubrey R. Odom-Mabey, W. Evan Johnson, Jason H. Yang, A. Murat Eren, Roland Brosch, Pradeep Kumar, David Alland

**SUPPLEMENTARY MATERIAL**

As part of Bact-Builder development and performance testing of individual steps, we generated simulated long and short reads off the H37Rv (NC_000962.3) linear reference genome. data using BadReads[26] and ART [27], respectively. These reads provided us with consistent high quality input data for initial validation and comparative analysis of robustness and reproducibility among various optional tools in the pipeline, allowing us to remove common experimental variables that can affect sequencing output and quality (ex. gDNA and sequencing library quality). We compared four commonly used long read assembly tools: Canu, Flye, Miniasm, and Raven, used in triplicate to assemble *in silico*-generated long read sequences. We found that each assembler produced a single linear contig, although the contig size varied across different assemblers (Figure S1A,B). This contig was not circularized as the *in silico* reads were generated off a linear genome. Next, in order to determine the degree of sequencing coverage required for optimal assemblies, our *in silico* data was subsetted using Rasusa [70] into individual read sets containing 30x, 50x, 100x, 250x, and 500x coverage. Previous studies have demonstrated that bacterial genomes can be fully assembled using as little as 30x long read sequencing coverage [18,71,72], however bacteria with high GC content often require higher coverage [71]. Testing our assembler against *in silco* generated reads, we determined that coverage higher than 30x did not lead to significant improvements in long read assemblies (Figure S1B).

Further comparative analysis of assembler differences using anvi'o [38] revealed that these size differences did not substantially change annotations across any of the individual assemblies, apart from Miniasm which was missing 2 gene clusters observed in other assemblies. However, a further comparison between individual assemblers using DNAdiff [36] revealed differences in SNP and indel counts and several regions of difference when compared to the H37Rv1998 reference, even though the reads were created using the same reference.

This *in silico* data analysis revealed inherent assembly differences associated with specific assemblers, suggesting the need for a better standard assembly tool that would consistently and correctly reconstruct microbial genomes. We attempted to reconcile the differences observed across assemblers using Trycycler[52] to generate a consensus sequence from the outputs of the four individual assemblers, and further improve it by polishing steps using both long- and short-reads. Using in *silico* generated long reads, Trycycler generated a consensus assembly that was 4,411,524 bases which was closer in size to the H37Rv reference (4,411,532 bp) than most individual assemblies (Table S3, Figure S1A,B). We next evaluated the optimal approach for final polishing steps, examining several iterations of both long and short read polishing tools (Table S2). An evaluation of polishing output, ease of use and amount of upstream data manipulation was used to determine the final workflow of three rounds of Racon [73] using long read data, followed by one round of Medaka (https://github.com/nanoporetech/medaka) with long read data and finally three rounds of Pilon [35] polishing using Illumina data. Nextflow logs detail statistics on CPU usage, memory, job duration and input/output for *in silico* analysis (Supplementary figure 2, Table 2). The *silico* data based assembly did not require basecalling or demultiplexing, and the remaining steps: assembly, Trycycler and polishing took approximately 15.8 hours on a standard CPU node to complete (Figure S2A-D).

| **Supplementary Table 1.** Library prep protocols for Illumina and ONT sequencing | | | |
|---|---|---|---|
| **Company** | **Kit Name** | **Minimum Input required** | **Source** |
| Illumina | PCR-free Tagmentation | 25-300 ng | https://www.Illumina.com/products/by-type/sequencing-kits/library-prep-kits/truseq-dna-pcr-free.html |
| ONT (MinION) | Ligation Sequencing Kit | 1000ng | https://store.nanoporetech.com/us/sample-prep/ligation-sequencing-kit.html |

**Supplementary Table 2.** Various polishing tools and combinations evaluated for Bact-Builder pipeline

| Name | # of contigs | Size (bp) | SNP* count relative to H37Rv reference | Indel** count relative to H37Rv reference |
|---|---|---|---|---|
| Racon (Illumina)x3 + Medaka (ONT) + Pilon (Illumina)x3 | 1 | 4411530 | 0 | 2 |
| Racon (Illumina)x3 + Medaka (ONT) | 1 | 4411402 | 0 | 134 |
| Racon (ONT)x3 + Medaka (ONT) + Pilon (Illumina)x3 | 1 | 4411530 | 0 | 2 |
| Medaka (ONT) + Pilon (Illumina)x3 | 1 | 4411530 | 0 | 2 |

*SNP: Single Nucleotide Polymorphism; ** Indels: single base insertions or deletions

**Supplementary table 3.** *In silico* assembly results and DNAdiff analysis results comparing individual assemblies to the H37Rv reference (H37Rv ref)

| **Name** | **# of contigs** | **Size (bp)** | **SNP* count (relative to ref)** | **Indel** count (relative to ref)** | **Regions of difference (relative to ref)** |
|---|---|---|---|---|---|
| H37Rv ref | 1 | 4411532 | - | - | - |
| H37Rv *in silico* canu1 | 1 | 4411506 | 0 | 0 | 2 |
| H37Rv *in silico* canu2 | 1 | 4411507 | 0 | 0 | 2 |
| H37Rv *in silico* canu3 | 1 | 4411270 | 0 | 0 | 2 |
| H37Rv *in silico* flye1 | 1 | 4411531 | 0 | 0 | 1 |
| H37Rv *in silico* flye2 | 1 | 4411531 | 0 | 0 | 1 |
| H37Rv *in silico* flye3 | 1 | 4411531 | 0 | 0 | 1 |
| H37Rv *in silico* miniasm 1 | 1 | 4411567 | 4 | 118 | 1 |
| H37Rv *in silico* miniasm 2 | 1 | 4411533 | 2 | 101 | 2 |
| H37Rv *in silico* miniasm 3 | 1 | 4411564 | 10 | 114 | 2 |
| H37Rv *in silico* raven1 | 1 | 4411460 | 1 | 4 | 2 |
| H37Rv *in silico* raven2 | 1 | 4411394 | 0 | 5 | 2 |
| H37Rv *in silico* raven3 | 1 | 4411465 | 0 | 4 | 2 |
| H37Rv *in silico* Trycycler | 1 | 4411524 | 0 | 0 | 2 |
| H37Rv *in silico* Trycycler + Polish | 1 | 4411524 | 0 | 0 | 2 |

*SNP: Single Nucleotide Polymorphism; ** Indels: single base insertions or deletions

### Supplementary Figure 1

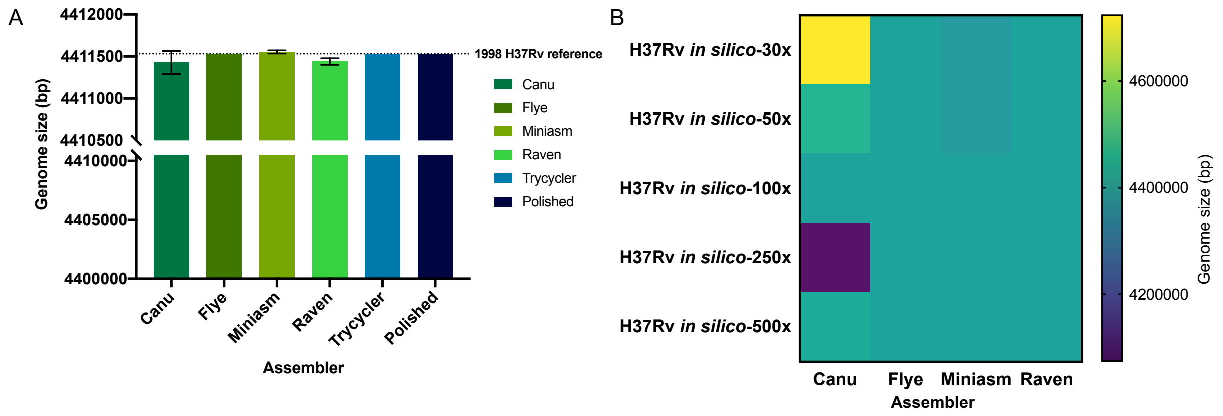


**Supplementary figure 1. Developing Bact-Builder A.** Comparison of *in silico* assembly sizes generated by each assembler. **B.** Heatmap comparison of genome sizes of the four de novo long read assemblers of using *in silico* generated sequencing reads. The sequence coverage sampled for each analysis is shown in each row on the Y axis.

### Supplementary Figure 2

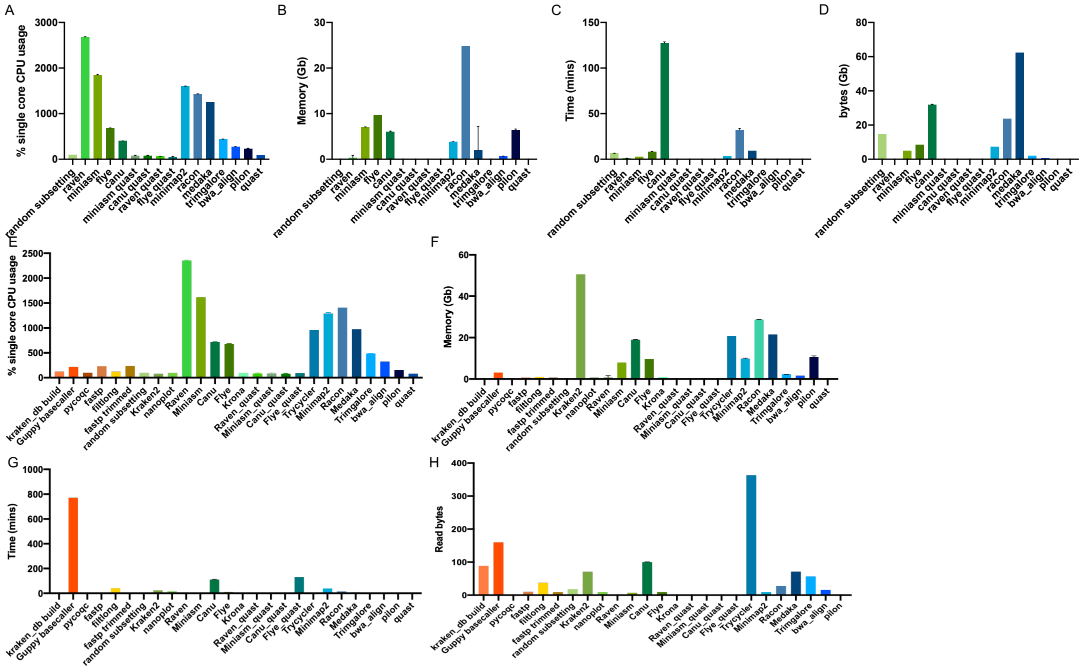


**Supplementary Figure 2. Summary information provided by Nextflow output for assembly of H37Rv *in silico* data and H37Rv.1 comparing all 4 tested assemblers and polishing. A-D.** H37Rv *in silico* data **A.** Amount of CPU usage **B.** Memory (RAM) usage **C.** Time to completion *D.* Input/Output (I/O) (how much data is read per task). **E-H** H37Rv.1 data. **E.** Amount of CPU usage **F.** Memory (RAM) usage **G.** Time to completion H. Input/Output (I/O) (how much data is read per task).

### Supplementary Figure 3

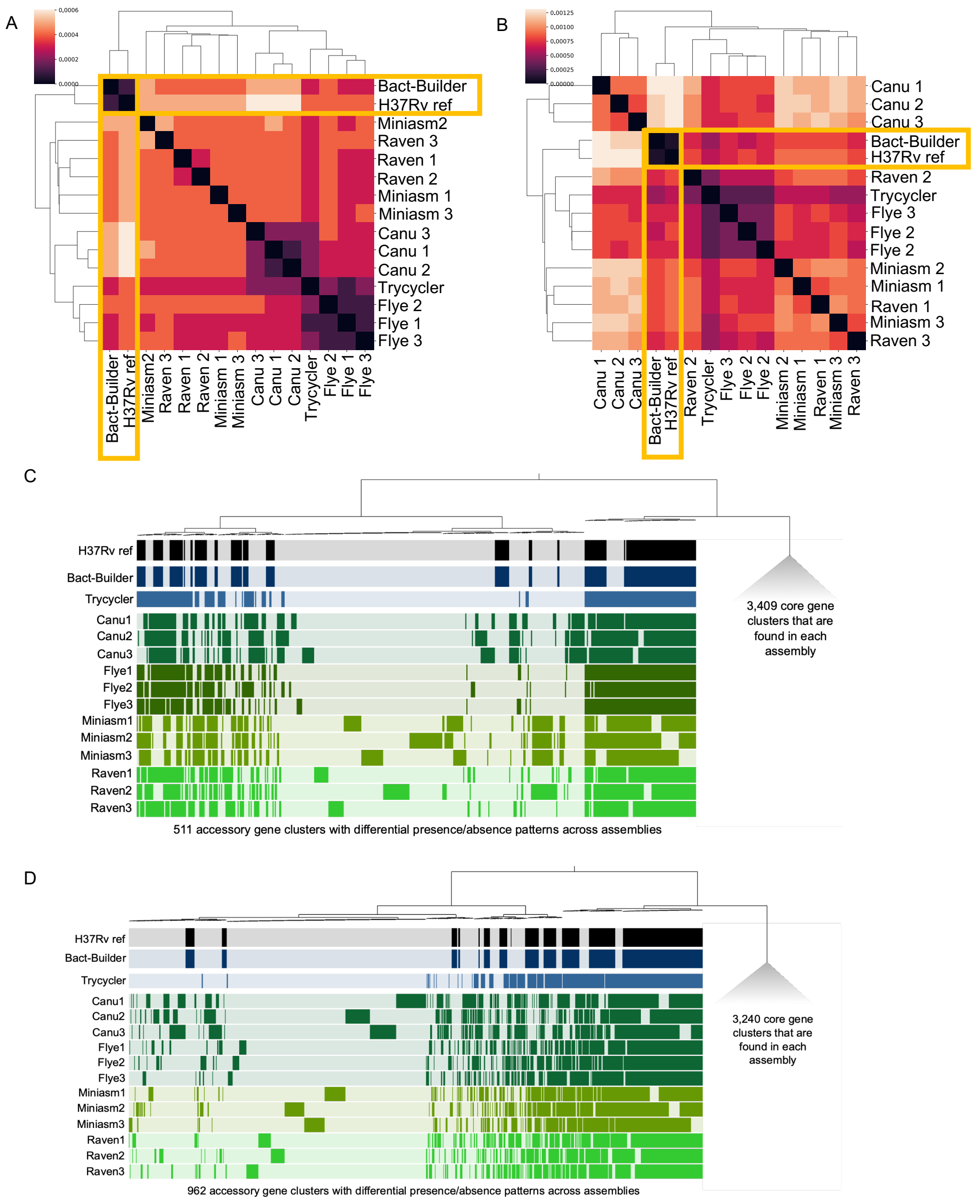


**Supplementary figure 3. Comparing H37Rv.2 and H37Rv.3 assemblies**. **A-B.** Heatmap of hierarchical clustering of the distance using euclidean average linkage clustering of differences between all assemblies for H37Rv.1(**A**) and H37Rv.3(**B**), the Bact-Builder output and the published reference (H37Rv ref) determined by DNAdiff. **C-D**. Anvio output showing differences in annotations between H37Rv.2(**C**) and H37Rv.3 (**D**).

**Supplementary Figure. 4**

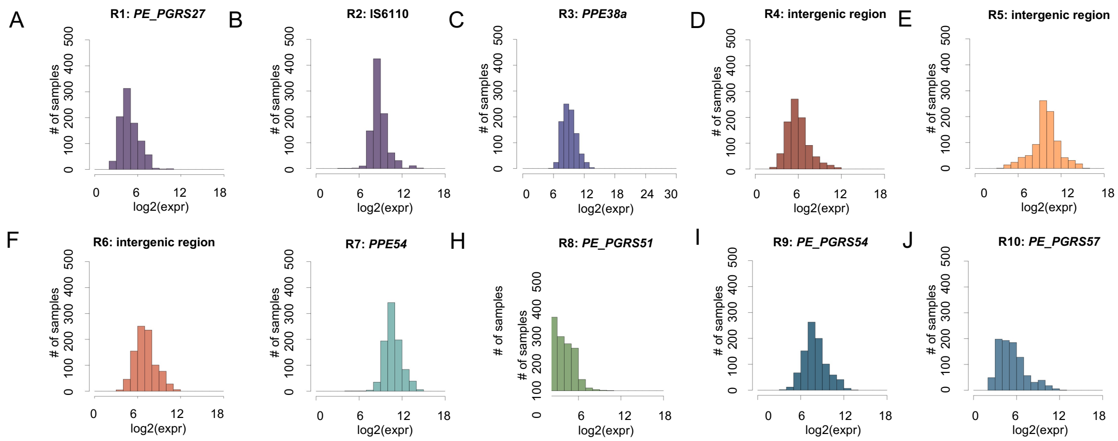


**Supplementary figure 4. RNA counts of newly discovered genomic regions of difference from public RNA sequencing datasets.** (A-J) Histogram of R1 – R10 respectively. Histograms demonstrate that all regions are expressed in H37Rv (new).
